## Supplementary figures and images for "*Drosophila* models of pathogenic copy-number variant genes show global and non-neuronal defects during development"

### S1 Figure

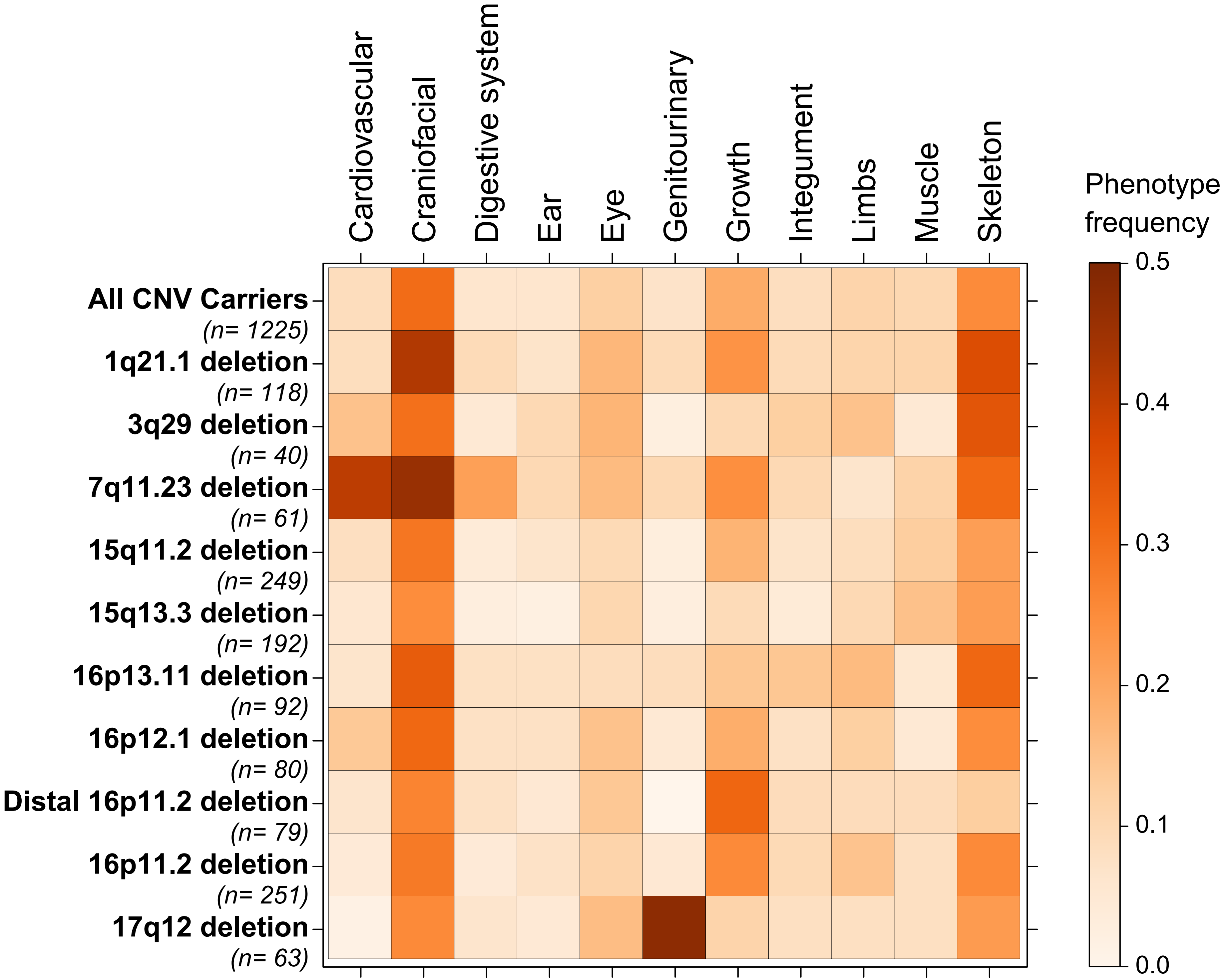

### S2 Figure

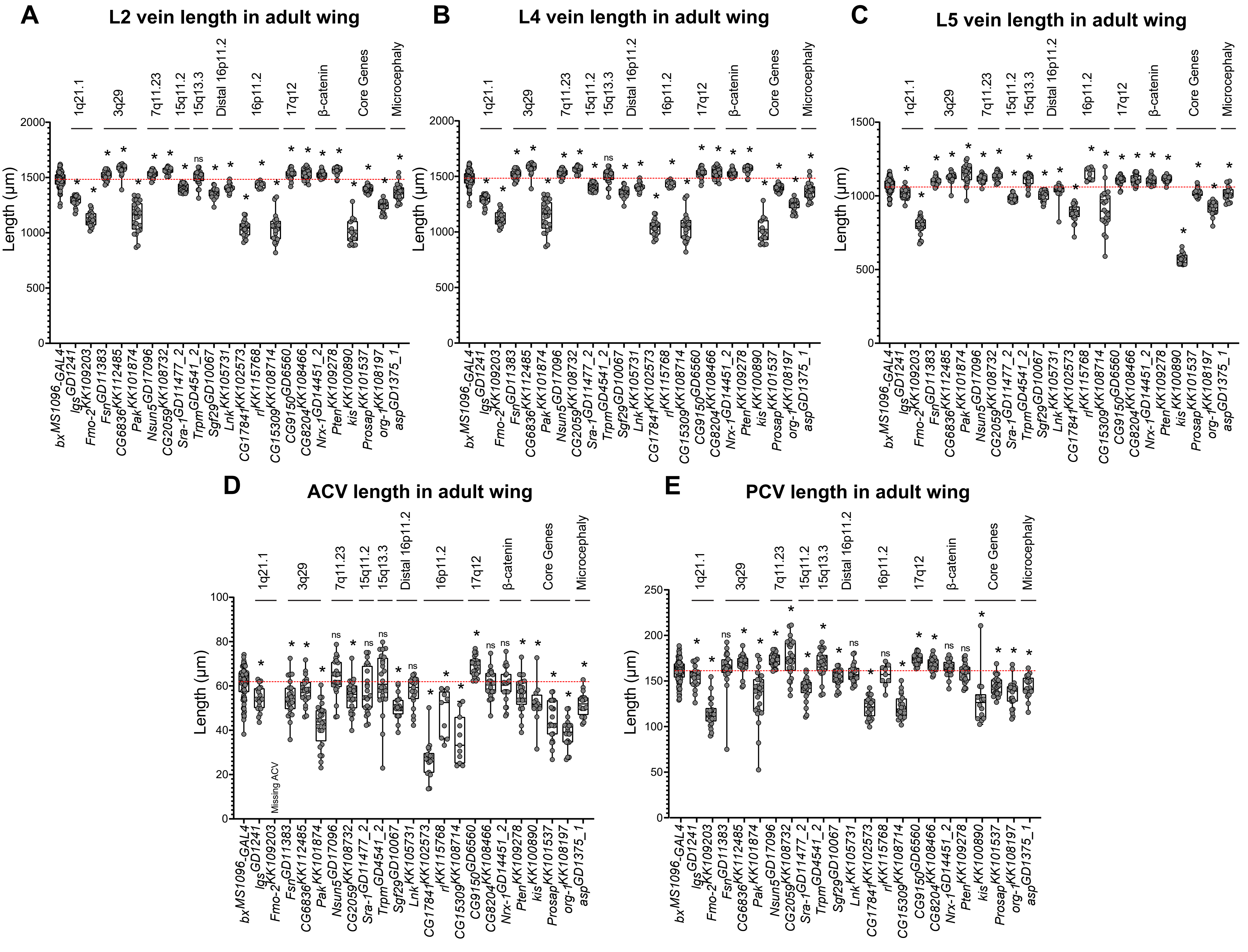

### S3 Figure

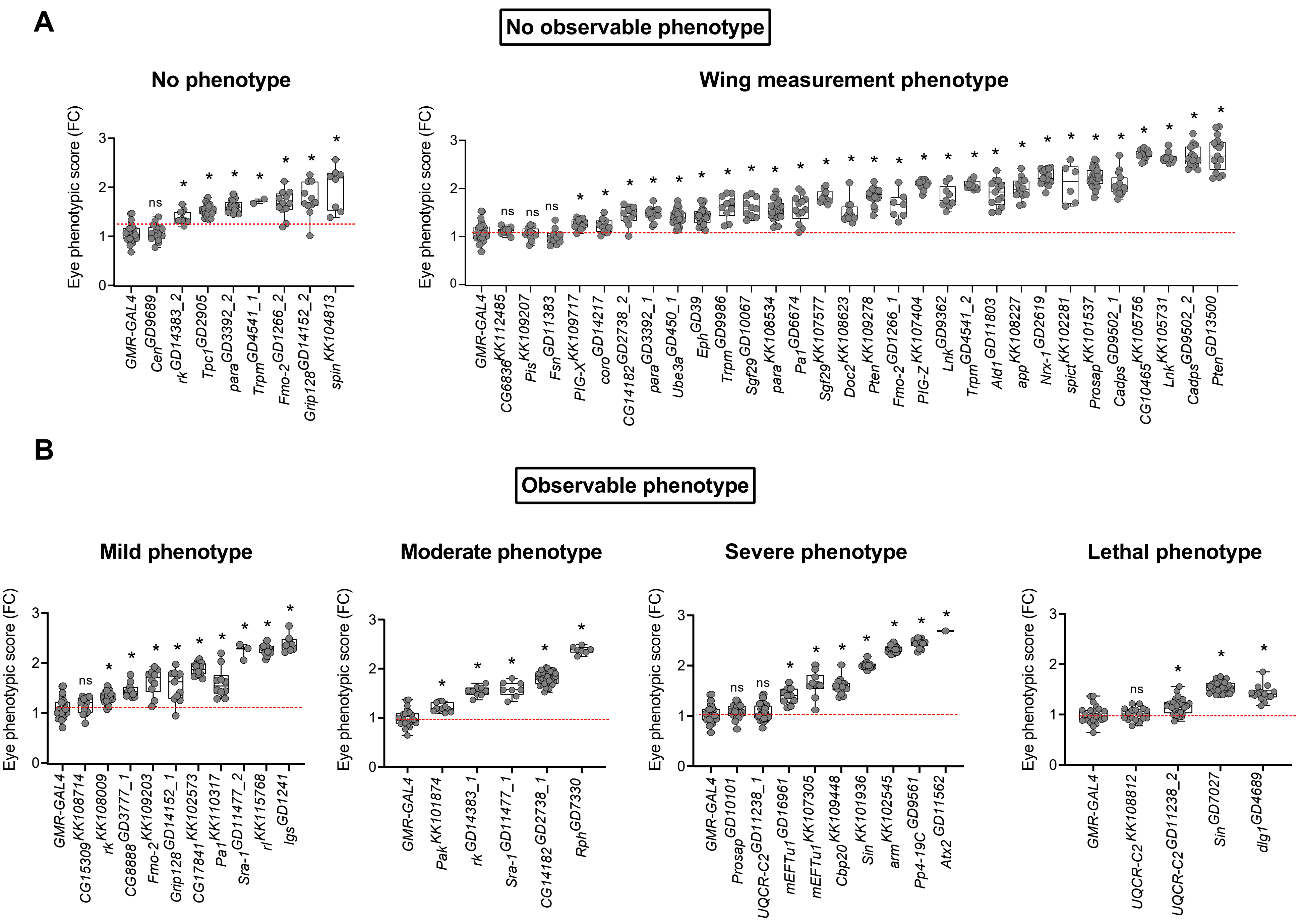

### S4 Figure

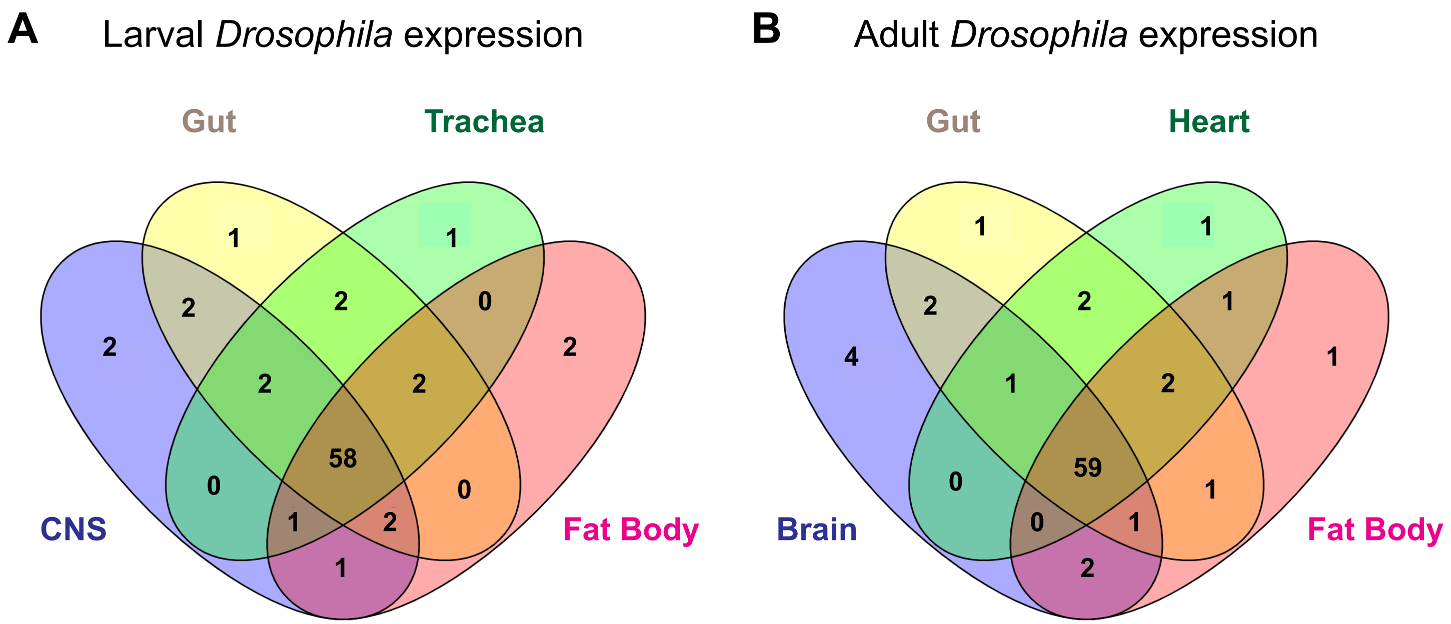

### S5 Figure

#### Cellular processes in larval wing discs

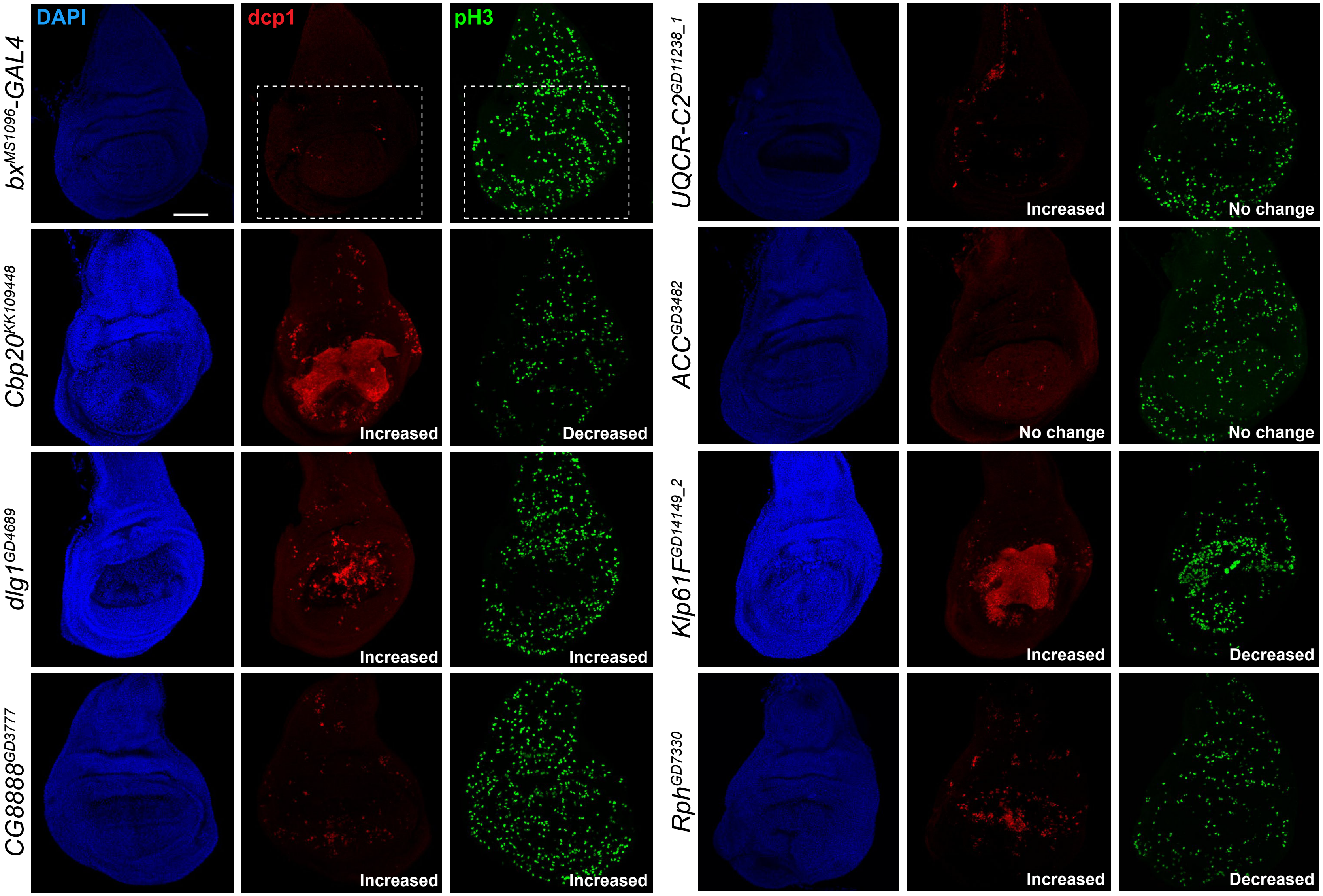

### S6 Figure

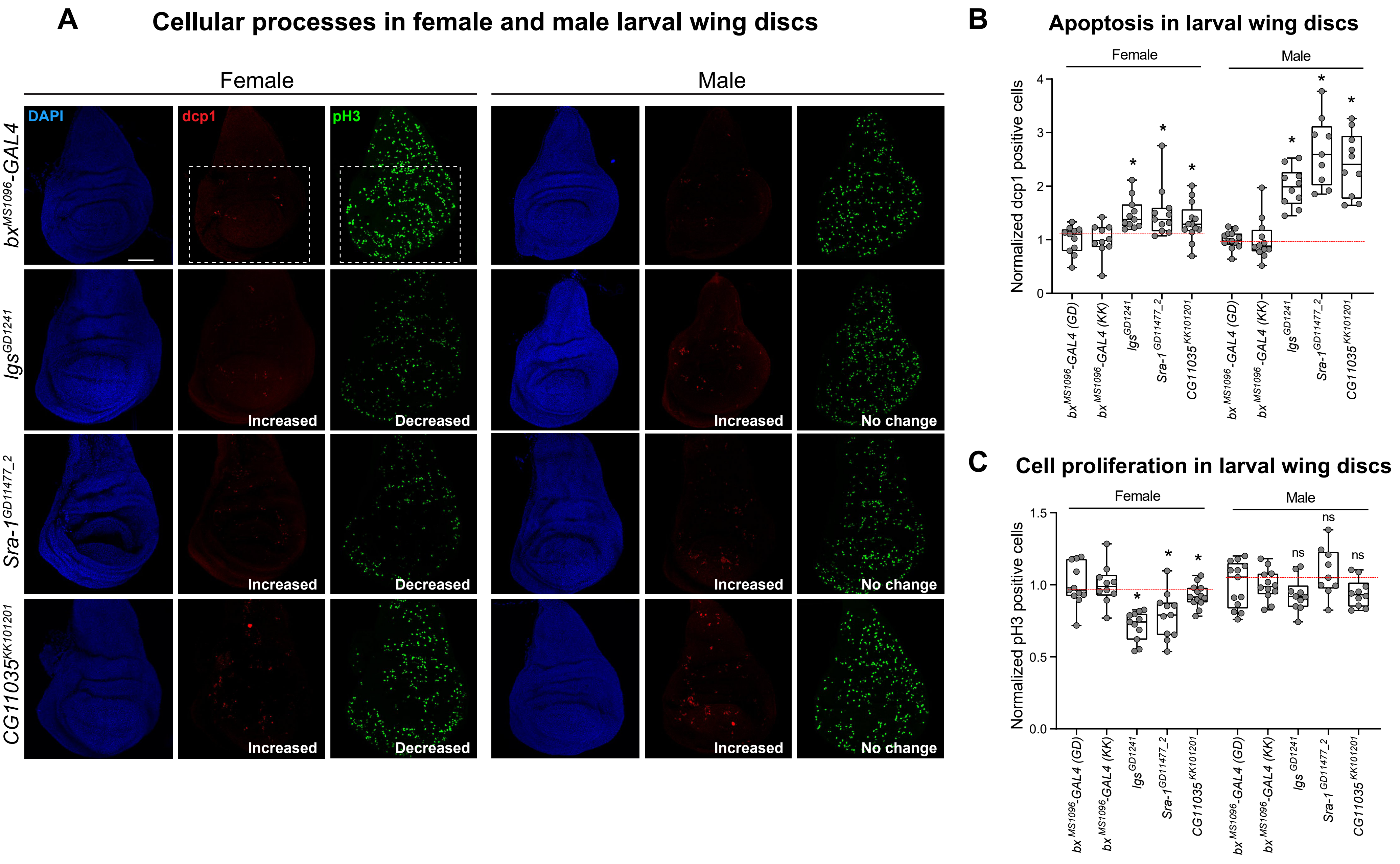

### S7 Figure

#### Disruption of signaling pathways in larval wing discs

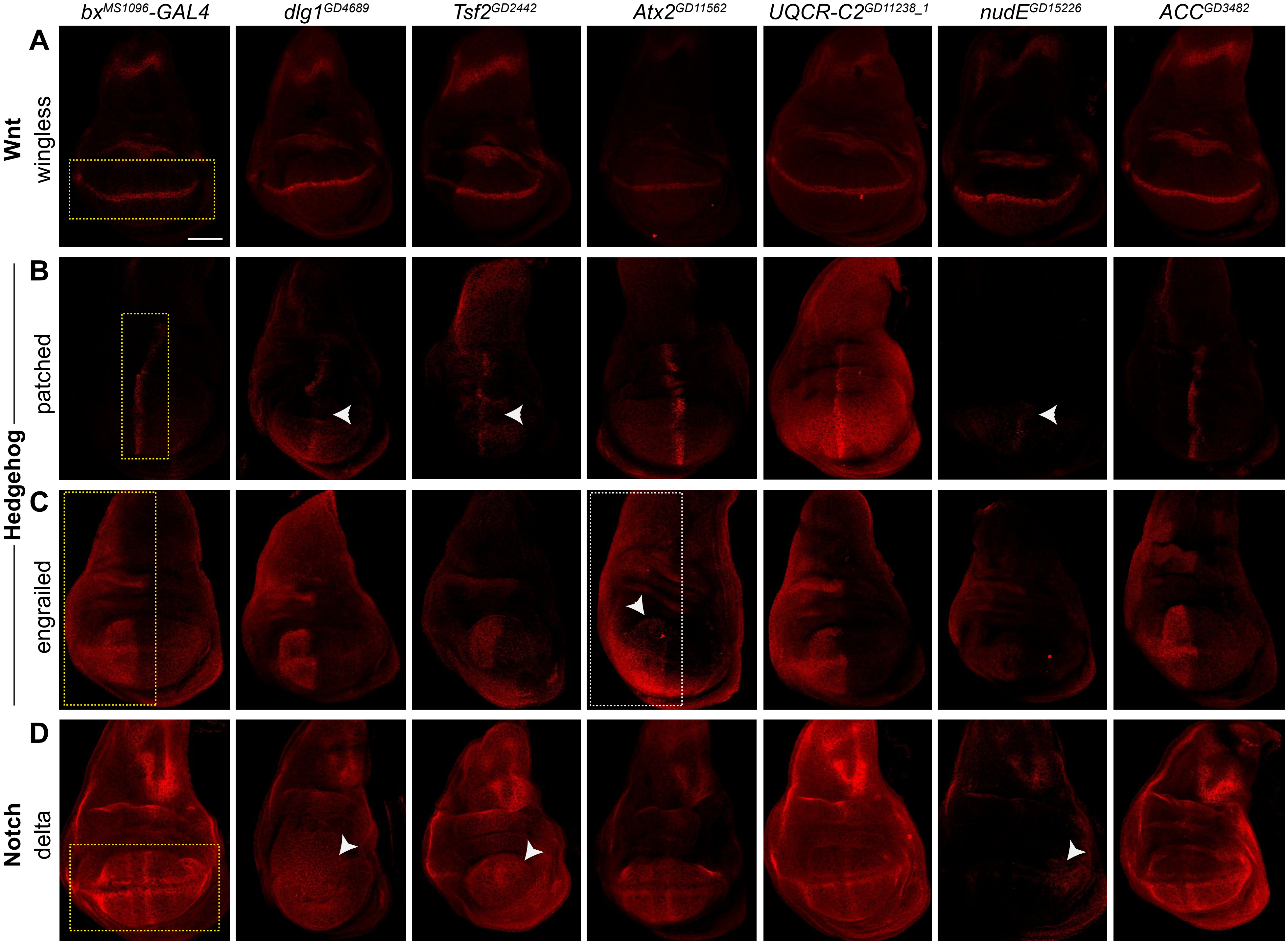

### S8 Figure

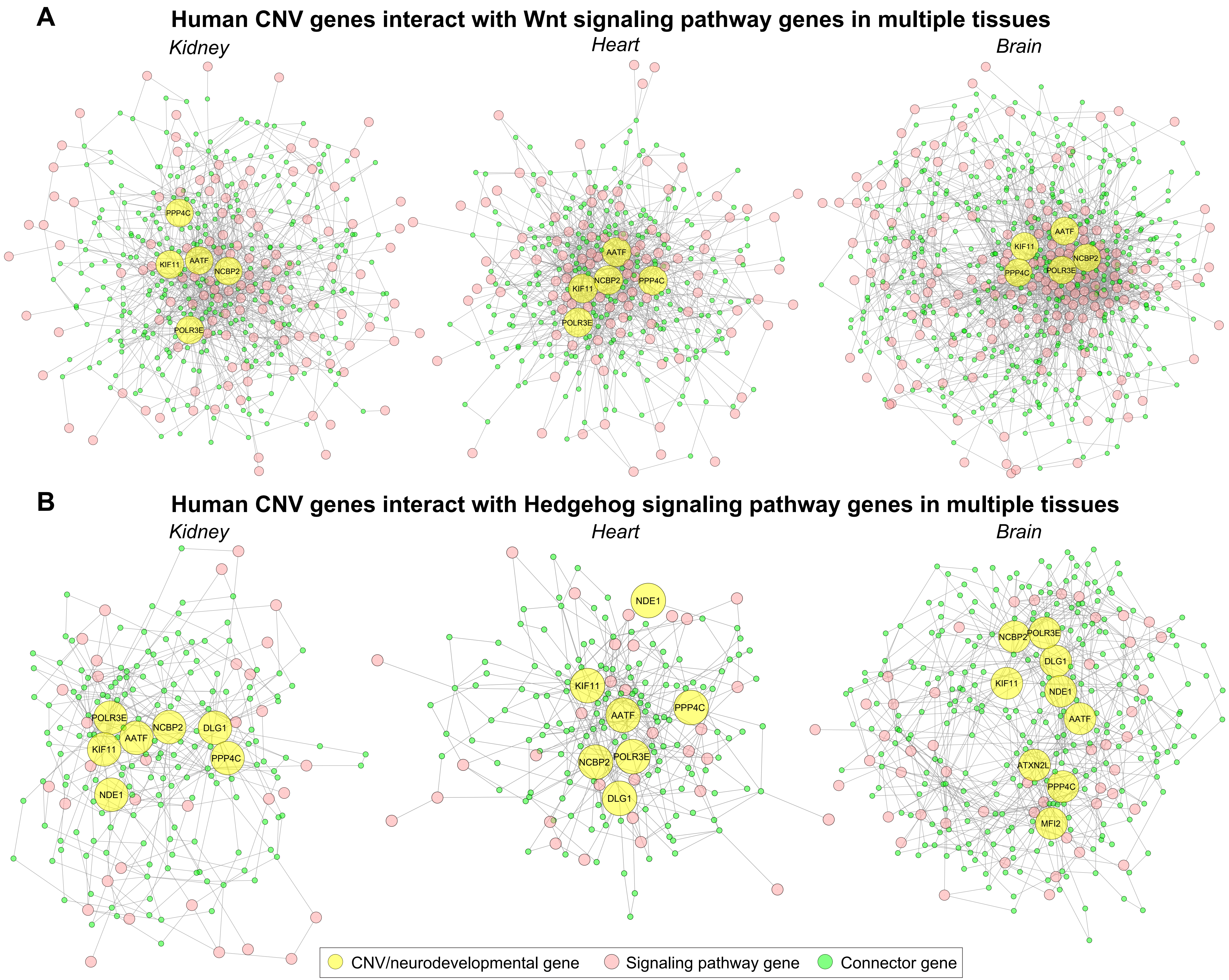
